# Divergent binding mode for a protozoan BRC repeat to RAD51

**DOI:** 10.1101/2022.01.25.477309

**Authors:** Teodors Pantelejevs, Marko Hyvönen

## Abstract

Interaction of BRCA2 through ca. 30 amino acid residue motifs, BRC repeats, with RAD51 is a conserved feature of the double-strand DNA break repair process in eukaryotes. In humans the binding of the eight BRC repeats is relatively well understood, with structure of BRC4 repeat bound to human RAD51 showing how two sequence motifs, FxxA and LFDE, in the BRC repeat interact with distinct sites on RAD51. Little is known however of the interaction of BRC repeats in other species, especially in protozoans where variable number of BRC repeats are found in BRCA2 proteins. Here we have studied in detail the interactions of the two BRC repeats in *Leishmania infantum* BRCA2 with RAD51. We show that the *Li*BRC1 is a high affinity repeat with a K_D_ of 0.29 μM while *Li*BRC2 binds to RAD51 with a K_D_ of 13.5 μM. A crystal structure of *Li*BRC1 complexed with *Li*RAD51 revels an extended β-hairpin compared to human BRC4 and shows that the equivalent of human LFDE motif is not interacting with *Li*RAD51. A truncation analysis of *Li*BRC1 confirms that a shorter repeat is sufficient for high affinity interaction and this minimal repeat is functional in inhibiting the formation of *Li*RAD51-ssDNA nucleofilament.

## INTRODUCTION

Genomic integrity is critical for the survival of all forms of life, and successful repair of DNA lesions is an essential function of the cell. Eukaryotes have evolved a sensitive and highly organised response to DNA damage, which senses genotoxic events and elicits an appropriate repair cascade.^1^ Double-strand breaks (DSBs) are the most severe type of genotoxic damage that can result in irreversible genomic rearrangements, aneuploidy and cell death.^1^ In eukaryotes, DSB repair can happen via several mechanisms, namely, homologous recombination (HR), non-homologous end joining (NHEJ),^2^ microhomology-mediated end joining (MMEJ)^3^ and single strand annealing (SSA).^4^

HR is the most faithful DSB repair pathway, as it employs a DNA template that is homologous to the broken locus in order to resynthesise the DNA at the double-strand break, and thus restores the original nucleotide sequence.^5^ This is in contrast to NHEJ, MMEJ and SSA, which can introduce small but potentially detrimental changes to the genome.^6,^ A mitotic sister chromatid is the preferred DNA donor template for HR, but repair can also proceed using the corresponding homologous chromosome or other homologous loci in the genome, which can lead to the loss of heterozygosity.^7,8^ Due to the requirement for a sister chromatid, HR happens predominantly during S and G2 phases of cell cycle.^9^

The RAD51 recombinase is the central mediator of mitotic homologous recombination.^10^ HR is initiated by the resection of the 5′ strand at an end of a DSB, resulting in a 3′ ssDNA overhang.^11,12^ Oligomeric RAD51 binds to the resected strand, forming a pre-synaptic nucleofilament (NF), which then invades homologous dsDNA that serves as the template for repairing the lesion.^13,14^

The human tumour suppressor BRCA2 is the most well-known regulator of RAD51, manifesting stimulatory effects on its function.^15^ Two distinct RAD51-binding regions have been identified in human BRCA2. The C-terminal TR2 region has a role in stabilising the RAD51:ssDNA nucleofilament.^16,17^ In the central part of BRCA2, encoded by the exon 11 in humans, are located a series of eight evolutionarily conserved ∼30-40 residue long sequence fragments termed BRC repeats that are critical in regulation of RAD51 function.^18^

The first structure of a BRC repeat in complex with human RAD51 was determined by Pellegrini and colleagues using X-ray crystallography.^19^ The model shows that BRC repeat 4 (BRC4) binds the ATPase domain of RAD51 and reveals a number of critical structural features, or “hot-spots”, that drive the interaction. The most outstanding feature of the binding mode is the interface formed by the conserved FxxA motif (FHTA in BRC4, residues 1524-1527), which interacts with RAD51 at the FxxA site where an analogous motif in RAD51 mediates its self-association. While essential for BRC repeat binding to RAD51, this short motif alone is not sufficient to mediate high affinity interaction between the two proteins.^20,21^ At its C-terminal half, spanning residues Lys1536 to Glu1548, BRC4 folds into an α-helix that produces additional contacts with the ATPase domain through a combination of hydrophobic and polar interactions. Residues Ile1534, Leu1539, Val1542, Leu1545 and Phe1546 form a continuous hydrophobic interface with RAD51 by projecting their side chains into the ATPase domain surface. Two of these conserved hydrophobic residues, Leu1545 and Phe1546, bind a pronounced cognate pocket on the ATPase domain, and are followed by a conserved acidic Glu1548, which interacts with nearby arginine side-chains. This “LFDE” motif has been shown to be critical for high-affinity binding *in vitro* and is required for RAD51 function in cells.^20^

Here, we investigated the interaction between RAD51 and BRCA2 orthologs in *Leishmania infantum*, a protozoan parasite that harbours only two BRC repeats in its functional BRCA2 ortholog.^22^ We use biochemical and biophysical methods to evaluate the affinities of *L. infantum (Li)* BRC repeats to *Li*RAD51, define the minimal repeat that is needed for this interaction and characterise the complex of the higher affinity repeat with *Li*RAD51 by X-ray crystallography.

## MATERIALS AND METHODS

### Reagents and biological resources

All oligonucleotides used for cloning were obtained from Sigma-Aldrich and are provided in **Table S2**.

| Reagent | Source | Identifier |
| --- | --- | --- |
| **T7 Express *E. coli* cells** | New England Biolabs | C2566I |
| **BL21(DE3) *E. coli* cells** | New England Biolabs | C2530H |
| **FAM-dT60 oligonucleotide** | Sigma Aldrich | n/a |
| **TEV protease** | Prepared in house using the pRK793 vector | Addgene #8827 |
| **Fluorescein-5-Maleimide** | ThermoFisher | 62245 |
| ***Li*RAD51 synthetic gene** | GeneArt (Thermofisher), sequence provided in Supplementary Data. |  |
| **pPEPT1 plasmid** | Dr Teodors Pantelejevs, unpublished |  |
| **pEXP-MBP plasmid** | Dr Aleksei Lulla, unpublished | Addgene #112568 |
| **pHAT2 plasmid** | Dr Marko Hyvönen | Addgene #112583 |
| **pBAT4 plasmid** | Dr Marko Hyvönen | Addgene #112580 |

### Expression plasmid preparation

All protein expression constructs were cloned using sequence and ligation independent cloning (SLIC) using the primers provided in **Table S2**. DNA encoding the full-length *Li*RAD51 protein was codon-optimised for expression in E. coli and obtained as a synthetic gene from GeneArt (Thermofisher). Full-length *Li*RAD51 was cloned into pExp-MBP plasmid (Dr Aleksei Lulla, unpublished, Addgene #112568), as fusion to a TEV-cleavable maltose-binding protein expression tag. *Li*RAD51^ATPase^ (residues 134-386) was cloned into the pHAT2 vector (Dr Marko Hyvönen, unpublished, Addgene #112583), fused to an N-terminal His-tag. The DNA insert for *Li*RAD51^ATPase,ΔL2^ was prepared by removing residues 309-324 from *Li*RAD51^ATPase^ by overlap extension mutagenesis and cloned into a pBAT4 vector lacking any fusion tags (Dr Marko Hyvönen, Addgene #112580).

For *Li*BRC repeat expression constructs, the repeat DNA was first codon optimised for *E. coli* expression and oligonucleotides were designed using the DNAworks application.^23^ DNA inserts were prepared by assembly PCR and cloned into the pPEPT1 vector (Dr Teodors Pantelejevs, unpublished), containing an N-terminal GB1 tag and a C-terminal His_8_-Tag, or the pOP3BT vector (Dr Marko Hyvönen, unpublished, Addgene #112603), containing an N-terminal, TEV-cleavable His_8_-GB1 fusion.

### Expression and purification of proteins

All expression vectors were transformed into either T7Express (New England Biolabs) or BL21(DE3) *E. coli* cells and stored as glycerol stocks. For all protein constructs, cells were plated on LB agar supplemented with ampicillin (100 μg/ml) and grown overnight at 37 °C. Next day, cells were scraped and used to inoculate flasks containing 1 L of 2x YT medium supplemented with 100 μg/mL ampicillin. Cultures were grown at 37 °C until OD_600_ of 0.5-1.0, after which temperature was adjusted depending on the protein being expressed. After expression, prior cell lysis, cells from all constructs were supplemented with DNase I (100 μL, 2 mg/mL) and AEBSF (1 mM), and lysed on an Emulsiflex C5 homogenizer (Avestin) or by sonication. Cell lysate was centrifuged at 40 000 x*g* for 30 min and supernatant collected. Specific expression and purification steps are described for each individual construct. After the final purification step, proteins were concentrated to 0.5-1 mM in size exclusion chromatography (SEC) buffers and flash-frozen.

### Purification of monomeric *Li*RAD51^ATPase^

Expression was induced at 15 °C with IPTG (400 μM) overnight. Next day, cells were resuspended in 25 mL of IMAC buffer A (50 mM Tris-HCl pH 8.0, 150 mM NaCl, 100 mM Li_2_SO_4_, 20 mM imidazole) and frozen. Following cell lysis, lysate was loaded on a 3 mL Ni-NTA agarose matrix (Cube Biotech), after which column matrix was washed with 10 CV Nickel Buffer A. *Li*RAD51^ATPase^ was eluted with 12 ml IMAC buffer B (50 mM Tris-HCl pH 8.0, 150 mM NaCl, 100 mM Li_2_SO_4_, 200 mM imidazole). Protein was concentrated to 2 ml on a centrifugal filter (Amicon, MWCO 10 000 Da) and purified on a Superdex 75 16/60 HiLoad size exclusion column (Cytiva) equilibrated with 20 mM Tris pH 8.0, 100 mM NaCl, 100 mM Li_2_SO_4_.

### Purification of the *Li*BRC1: *Li*RAD51^ATPase,ΔL2^ complex

Separate flasks were inoculated with the GB1-*Li*BRC1 and *Li*RAD51^ATPase,ΔL2^ construct-expressing cells. Expression was induced with IPTG (400 μM) for three hours at 37 °C for the *Li*BRC1 cultures and overnight at 16 °C for the *Li*RAD51^ATPase,ΔL2^ cells. After expression, cells were resuspended in 25 mL of IMAC buffer A (50 mM Tris-HCl pH 8.0, 150 mM NaCl, 20 mM imidazole) and lysed. Lysate containing the GB1-*Li*BRC1 fusion was loaded on a 3 mL Ni-NTA agarose matrix (Cube Biotech), followed by the application of *Li*RAD51^ATPase,ΔL2^ lysate from equal culture volume. Column matrix was washed with 10 column volumes nickel buffer A. Complex was eluted with IMAC buffer B (50 mM Tris-HCl pH 8.0, 150 mM NaCl, 200 mM imidazole) into 2 ml fractions. Fractions containing the complex were pooled (∼10 ml total) and buffer-exchanged back into nickel buffer A on a PD-10 desalting column (Cytiva). Buffer-exchanged IMAC output was incubated with 100 μL of 2 mg/ml TEV protease overnight at 4°C. GB1 fusion partner was then removed from the solution by a second Ni-NTA affinity step, collecting the flow-through that contains the complex. Flow-through was concentrated on a centrifugal filter (Amicon, MWCO 3000 Da) to 2 ml and loaded onto a Superdex 75 16/60 HiLoad size exclusion column (Cytiva), previously equilibrated with SEC buffer (20 mM Tris pH 8.0, 100 mM NaCl, 100 mM Li_2_SO_4_, 1 mM EDTA). The complex eluted at ∼75 ml, fractions were analysed by SDS-PAGE.

### Purification of GB1-*Li*BRC-His_8_ fusions

Cells carrying pPEPT1 plasmids expressing GB1-*Li*BRC-His_8_ constructs were induced with IPTG (400 μM) for three hours at 37 °C. Cells were resuspended in 25 mL of IMAC buffer A (50 mM Tris-HCl pH 8.0, 150 mM NaCl, 20 mM imidazole) and frozen. After lysis and centrifugation, lysate was loaded on a 3 mL Ni-NTA agarose matrix (Cube Biotech), after which column matrix was washed with 10 column volumes IMAC Buffer A. Bound protein was eluted with 10 ml IMAC buffer B (50 mM Tris-HCl pH 8.0, 150 mM NaCl, 200 mM imidazole). The sample was diluted to 80 ml Q-A buffer (20 mM Tris-HCl, pH 8.0, 1 mM EDTA) and loaded on a HiTrap Q HP 5 ml column (Cytiva), which was washed with Q-A buffer, after which the GB1-*Li*BRC fusion was eluted with a linear 15 CV, 0-100% gradient of Q-B buffer (20 mM Tris-HCl, pH 8.0, 1 M NaCl, 1 mM EDTA).

### Preparation of the fluorescent polarisation probe

Cells carrying the pOP3BT-NCys-*Li*BRC1 plasmid were grown at 37 °C until OD_600_ of ∼1, after which expression was induced with IPTG (400 μM) for three hours. Cells were resuspended in 25 mL of IMAC buffer A (50 mM Tris-HCl pH 8.0, 150 mM NaCl, 20 mM imidazole, 0.5 mM TCEP). Lysate was loaded on a 3 mL Ni-NT A agarose matrix (Cube Biotech), after which column matrix was washed with 10 column volumes IMAC Buffer A. GB1-NCys-*Li*BRC1 was eluted with 12 ml IMAC buffer B (50 mM Tris-HCl pH 8.0, 150 mM NaCl, 200 mM imidazole, 0.5 mM TCEP). The eluent was buffer exchanged back into IMAC buffer A on a PD-10 desalting column (Cytiva). Buffer exchanged protein (∼18 ml) was incubated with 100 μL of 2 mg/ml TEV protease overnight at 4°C. The GB1 tag was then removed from the solution by a second IMAC step, collecting the flow-through that contains the NCys-*Li*BRC1 peptide. The flow-through was acidified with HCl to pH 2-4 and acetonitrile was added to 10%, after which the solution was centrifuged at 10000 x*g* for 15 min. The acidified flow-through was then applied to an ACE C8 300 4.6 × 250 mm semi-prep RP-HPLC column equilibrated with RPC buffer A (10% acetonitrile, 0.1% TFA) and peptide was eluted with a 20 CV gradient of RPC buffer B (90% acetonitrile, 0.1% TFA). Peak fractions were analysed by LCMS and pooled for drying under vacuum. The peptide was resuspended in PBS and labelled with Fluorescein-5-Maleimide (ThermoFisher), according to manufacturer’s instructions. This was followed by a second reversed phase chromatography step on an ACE C18 300 4.6 × 250 mm semi-prep RP-HPLC column, using identical buffers to the C8 step. Fluoresceinated peptide was dried and resuspended in MilliQ water. Mass of the peptide was confirmed by LCMS (**Figure S4**).

### Preparation of the *Li*BRC1 free peptide

*Li*BRC1 peptide was prepared in identical manner to the fluoresceinated *Li*BRC1 described above, except that no labelling and second reversed phase chromatography step was done.

### Isothermal titration calorimetry

*Li*BRC1 peptide was resuspended in MilliQ water to 10 times the desired concentration in the syringe. This was then diluted 10x with the ITC buffer (20 mM Tris pH 8.0, 150 mM NaCl, 100 mM Li_2_SO_4_) to obtain the final titrant solution. *Li*RAD51^ATPase^ was buffer-exchanged on a NAP-5 desalting column into ITC buffer and protein concentration was adjusted to 10:9 of the desired final value. One ninth volume of MilliQ water was added to the solution to bring the protein concentration to the desired final value, while maintaining identical buffer:MilliQ volume proportions in both the syringe and the cell. ITC was carried out using a Microcal ITC200 instrument at 25 °C with a 5.00 μCal reference power DP value, stirring speed of 750 rpm, 2 s filter period. ITC data were fitted using a single-site binding model using the Microcal ITC data analysis software in the Origin 7.0 package.

### Crystallography of the *Li*BRC1:*Li*RAD51^ATPase^ complex

Complex was diluted to a 0.5 mM concentration in SEC buffer (20 mM Tris pH 8.0, 100 mM NaCl, 100 mM Li_2_SO_4_, 1 mM EDTA). ADP/MgCl_2_ was added to the protein solution to a final concentration of 20 mM. Complex was crystallised in a 96-well MRC plate using the sitting-drop vapor diffusion technique. 200 nl of protein was added to 200 nl of precipitant containing 32% low MW PEG smear solution (Molecular Dimensions) and 100 mM Tris pH 8.5. Mosquito liquid handling robot (TTP Labtech) was used to dispense protein and reservoir solutions. Plates were stored at 17 °C in a RockImager crystallisation hotel (Formulatrix). Crystals were flash-frozen in liquid nitrogen without the addition of cryoprotectants. Diffraction data were collected on Diamond Light Source (Harwell, UK) beamline i04-1. Molecular replacement phasing method was used with the human RAD51 ATPase domain as search model (PDB: 1N0W). Molecular replacement was done with Phaser.^24^ The structure was refined without BRC repeats first and the peptides were built into the clearly visible electron density manually. Manual refinement was done in Coot and automated refinement with phenix.refine and autoBUSTER.^24,25^

### Fluorescence polarisation assay

All reactions were performed in black 384-well flat-bottom microplates (Corning) with a 40 μl final reaction volume. Following FP buffer was used: 50 mM Tris pH 8.0, 150 mM NaCl, 100 mM Li_2_SO4, 1% BSA, 0.1% Tween-20. Each reaction contained 5 nM of Fluor-NCys-*Li*BRC1 probe. For the direct titration experiment, *Li*RAD51^ATPase^ was added in two-fold serial dilutions. For competition experiments, *Li*RAD51^ATPase^ had a constant concentration of 500 nM and GB1-*Li*BRC repeats were added in serial dilutions instead. FP measurements were performed on a Pherastar FX (BMG Labtech) plate reader equipped with an FP 485-520-520 optic module. Each dilution was measured in triplicate. Graphs show means ± SD (n=3) per dilution. Binding curves were fitted using the four-parameter logistic model with a variable Hill slope using Prism software (Graphpad). Regression fitting was performed using the least squares optimisation algorithm. K_D_ values were estimated from the fitted IC_50_ parameters using a previously reported equation.^26^

### Electrophoretic mobility shift assay (EMSA)

*Li*RAD51:DNA-binding reactions (40 μl) were set up in 50 mM HEPES pH 7.4, 150 mM NaCl, 10 mM magnesium acetate, 2 mM CaCl_2_, 1 mM TCEP, 1 mM ATP. 5 μM *Li*RAD51 was incubated with varying concentrations of BRC repeats for 10 min at room temperature, followed by the addition of 100 nM fluorescently labelled FAM-dT60 oligonucleotide, and further incubation at 37 °C for 10 min. Control reactions were set up with free FAM-dT60 probe and FAM-dT60 + 5 μM *Li*RAD51. 10 μl of reactions were then loaded on a 1xTBE, 2% agarose gel and run at 250 V for 6 min at 4 °C. The gel was directly visualized on a Typhoon FLA 9000 imager (GE Healthcare) using FAM channels.

## RESULTS

### *Li*BRC1 binds *Li*RAD51 more strongly than *Li*BRC2

The interaction between the two *Li*BRC repeats and *Li*RAD51 was first qualitatively evaluated using affinity co-precipitation of the proteins from *E. coli* lysate. The *Li*BRC repeats (**Figure 1A)** were expressed containing an N-terminal G protein B1 domain (GB1 fusion), and a C-terminal His_8_-tag. A truncated, monomeric version of *Li*RAD51, containing only the ATPase domain (*Li*RAD51^ATPase^), was used to diminish competition for the FxxA site due to RAD51 self-oligomerisation. *Li*BRC1 co-precipitated *Li*RAD51^ATPase^ in near-stoichiometric amounts, suggesting a relatively strong interaction, whereas only trace amounts of *Li*RAD51^ATPase^ are seen in the *Li*BRC2 pull-down (**Figure 1B**).

**Figure 1.**
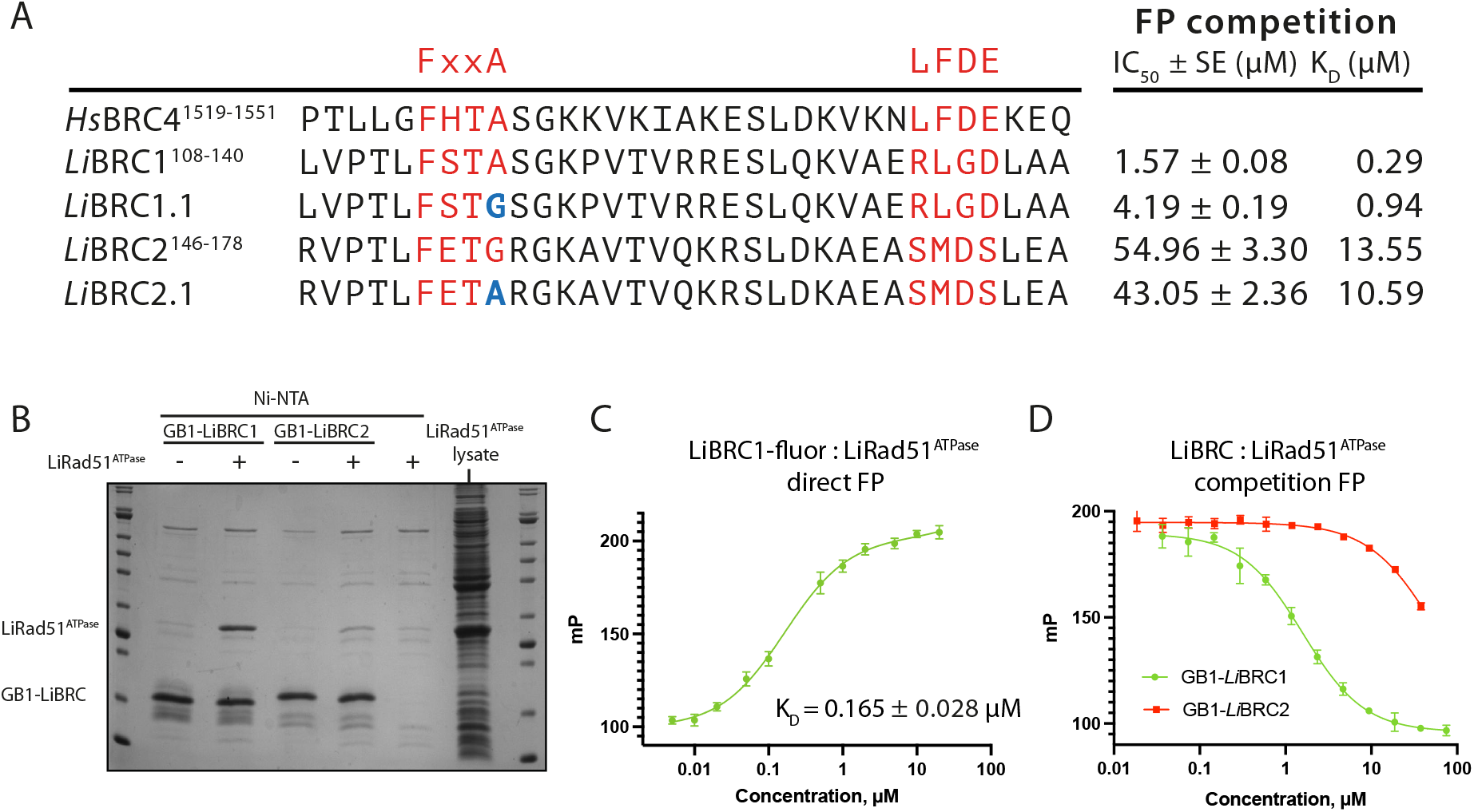
*Li*BRC1 is a more potent binder of *Li*RAD51 than *Li*BRC2. (**A**) Sequence alignment of the two *L. infantum* BRC repeats with *Hs*BRC4 and point mutants *Li*BRC1.1 and *Li*BRC2.1. Competition FP binding results are shown on the left. SE – standard error of fit. (**B**) Coomassie stained SDS-PAGE gel analysis of *L. infantum* BRC repeat affinity pull-down of *Li*RAD51^ATPase^ (**C**) Direct FP titration of *Li*RAD51^ATPase^ into fluorescein-tagged *Li*BRC1 (5 nM). Data shown are the means of triplicate measurements ± SE. (**D**) Competition FP titrations of GB1-fused *Li*BRC1 and *Li*BRC2. Fluorescently-labelled *Li*BRC1 probe (5 nM) was pre-incubated with 500 nM *Li*RAD51^ATPase^, to which GB1-*Li*BRC dilution series were added. Data shown are the means of triplicate measurements ± SD.

A fluorescence polarisation (FP) assay was used to evaluate the binding of the two repeats quantitively, using a fluorescent *Li*BRC1 peptide as a probe. Direct FP titration of *Li*RAD51^ATPase^ into this probe gave a K_D_ of 0.165 μM (**Figure 1C**). Competition experiments were then set up with purified GB1-fused peptide constructs, resulting in K_D_ values of 0.29 μM and 13.55 μM for GB1-*Li*BRC1 and GB1-*Li*BRC2, respectively (**Figure 1A,D**). To account for possible fusion partner-induced effects, *Li*BRC1 was also prepared as a free peptide and its K_D_ was determined by isothermal titration calorimetry (ITC) to be 0.65 μM, excluding the possibility that the GB1 tag has a significant effect on binding (**Figure S1**).

Previous studies on the human BRC repeats have shown that BRC5 is a low-affinity repeat, which has been rationalised by the mutation of an alanine to a serine in the FxxA motif of BRC5.^20,27^ We hypothesised that the affinity of *Li*BRC2 may be similarly diminished by a glycine instead of the alanine at the equivalent position. To test this, we mutated the *Li*BRC2 Gly154 to an alanine, however, this did not bring about a substantial increase in affinity, as determined by the competition FP assay (**Figure 1A**, *Li*BRC2.1, K_D_ = 10.59 μM). This observation further prompted us to investigate the role of the FxxA alanine in the context of the *Li*BRC repeats, and an FxxG mutant of *Li*BRC1 was likewise evaluated, displaying a three-fold reduction in affinity (**Figure 1A**, *Li*BRC1.1, K_D_ = 0.94 μM), similar to what has been observed for human FxxA tetrapeptide before.^21^ It is of note also that glycine is found in some archaeal RadA proteins instead of alanine in their self-association motif.^28^ It is thus reasonable to suggest that other factors besides the loss of a methyl group from the FxxA motif are responsible for the low affinity of *Li*BRC2.

The co-precipitation, FP and ITC data together show that *Li*BRC1 is a stronger *Li*RAD51 binder than LiBRC2 *in vitro*. Moreover, the affinity of both peptides is significantly lower than the nanomolar values reported previously for human BRC4 and other high-affinity repeats using similar monomeric forms of RAD51.^29,27^

### X-ray structure of the *Li*BRC1:*Li*RAD51 complex reveals novel binding features

In order to understand these interactions in more detail, we determined the crystal structure of the *Li*BRC1:*Li*RAD51^ATPase^ complex. To reduce the flexibility and conformational heterogeneity of the complex for crystallisation, a deletion mutant of the *Li*RAD51 ATPase domain was prepared by removing the DNA-binding loop L2 (residues 309-324), which in the absence of DNA is typically disordered and is not involved in BRC repeat binding (*Li*RAD51^ATPase,ΔL2^).

In the crystal structure, the overall fold of the *Li*BRC1 peptide is similar to what has been reported for *Hs*BRC4 (**Figure 2A**), with Phe113 and Ala116 hot-spot residues binding the two hydrophobic FxxA site pockets on the ATPase domain in an identical manner to BRC4. The β-turn, mediated by 115-TASGK-119, is also preserved and closely resembles the human BRC4 in its hydrogen-bonding pattern, with a Thr115 side-chain stabilising the turn through hydrogen bonding to Ser117 and to mainchain amide of Lys119 (**Figure 2B)**. Remarkably, an extended β-hairpin fold forms at the N-terminus of the repeat, similarly to what has been previously observed for the chimeric, high-affinity human BRC8-2 repeat (**Figure 2A,C)**.^27^ The β-hairpin is stabilised by a Thr111 residue, positioned -2 residues prior the FxxA motif, forming a hydrogen bonding network with the backbone amides of Val121 and Phe113 on the two anti-parallel strands of the repeat. The extended β-hairpin fold promotes the *Li*BRC1 peptide to form a small hydrophobic core mediated by the side-chains Val109, Thr111, Val123 and Leu128, as well as *Li*RAD51 surface residues Leu241, Gln242, Ala245, Met246 (**Figure 2C**). Leu112, which precedes Phe113, forms additional hydrophobic contacts with a hydrophobic cleft formed by *Li*RAD51 His235, Leu239 and Gln242. The sum of these observations suggest that *Li*BRC1 forms considerably more hydrophobic interactions at the FxxA site compared to human repeats with known structure.

**Figure 2.**
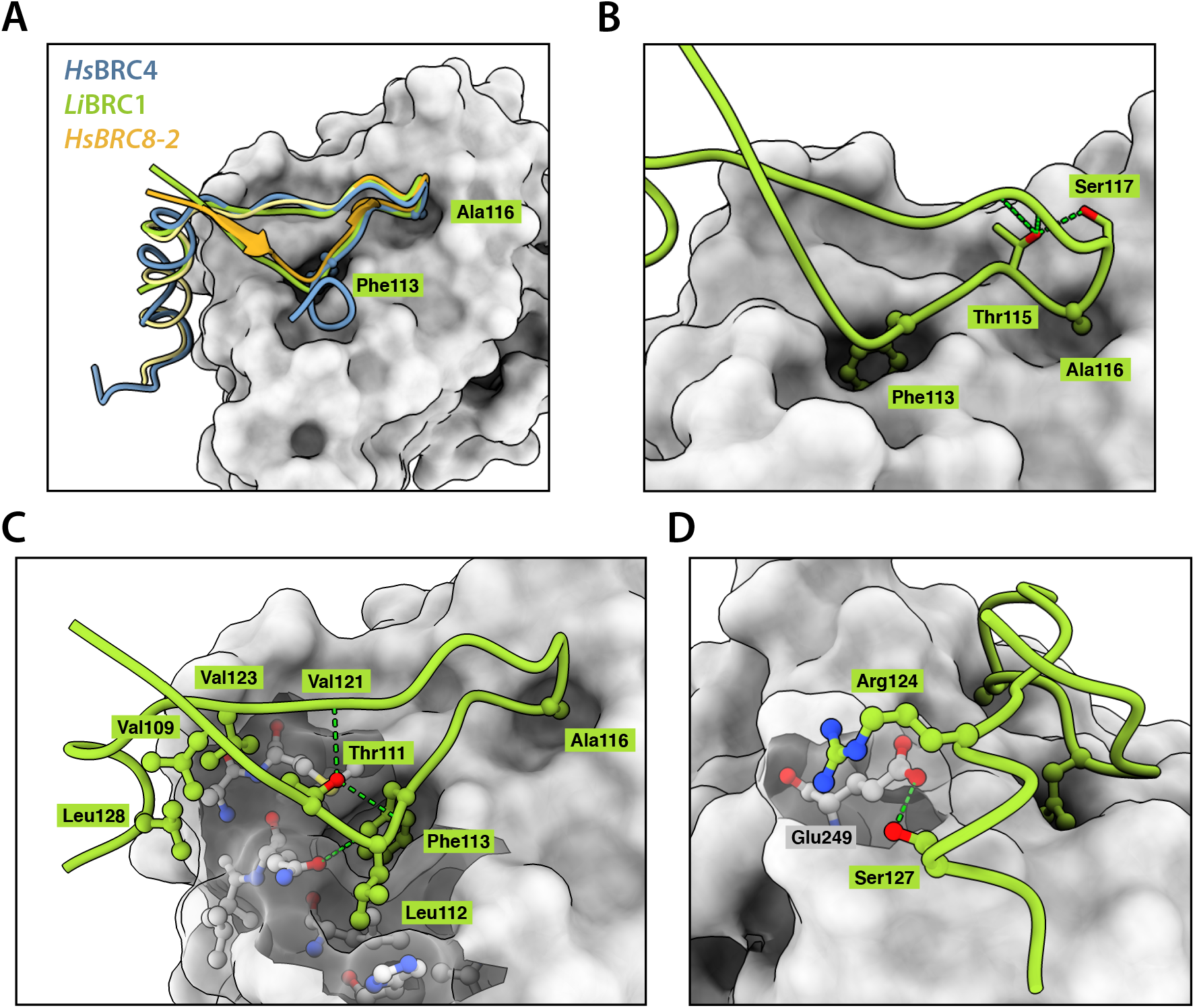
Crystal structure of *Li*BRC1 in complex with *Li*RAD51^ATPase^. (**A**) Overall binding mode of *Li*BRC1 (green) superposed with *Hs*BRC4 (blue) and *Hs*BRC8-2 (orange) (**B**) FxxA motif and the beta-turn mediated by Thr115 hydrogen-bonding (**C**) Detailed view of the extended β-hairpin fold and the hydrophobic interactions formed by *Li*BRC1 (**D**) Arg124-mediated electrostatic contacts with *Li*RAD51, further stabilised by a hydrogen bond with Ser127.

The C-terminal α-helix starts with a cationic Arg124 residue that forms electrostatic contacts with *Li*RAD51 Glu249 (**Figure 2D**). A similar interaction has not been observed for HsBRC4 or HsBRC8-2, despite human RAD51 also containing a glutamate at the equivalent position. Surprisingly, unlike the human BRC4 repeat, *Li*BRC1 lacks defined electron density beyond residue Gln129 (**Figure 3A,B**). In particular, the hot-spot residues of the LFDE motif, corresponding to RLGD in *Li*BRC1, have no discernible electron density even after the rest of the peptide has been modelled and several rounds of refinement done. To ensure that the C-terminus of the peptide was not degraded by bacterial proteases during purification, the complex was analysed by LCMS, and the full-length species of *Li*RAD51^ATPase,ΔL2^ and *Li*BRC1 were identified (**Figure S2**). The crystal structure suggests a binding mode for *Li*BRC1 in which the residues corresponding to the LFDE motif do not form critical contacts with the *Li*RAD51 ATPase domain. Comparison of the *Li*RAD51ATPase surface at this interface with that of the human RAD51 reveals structural features that support this binding mode. The *Li*RAD51 surface region corresponding to the pocket on human RAD51 where BRC4 Leu1545 and Phe1546 bind contains a significantly shallower cavity compared to human RAD51, resulting from Tyr205 and Met251 changing to Leu241 and Ser287, respectively (**Figure 3C-F**).

**Figure 3.**
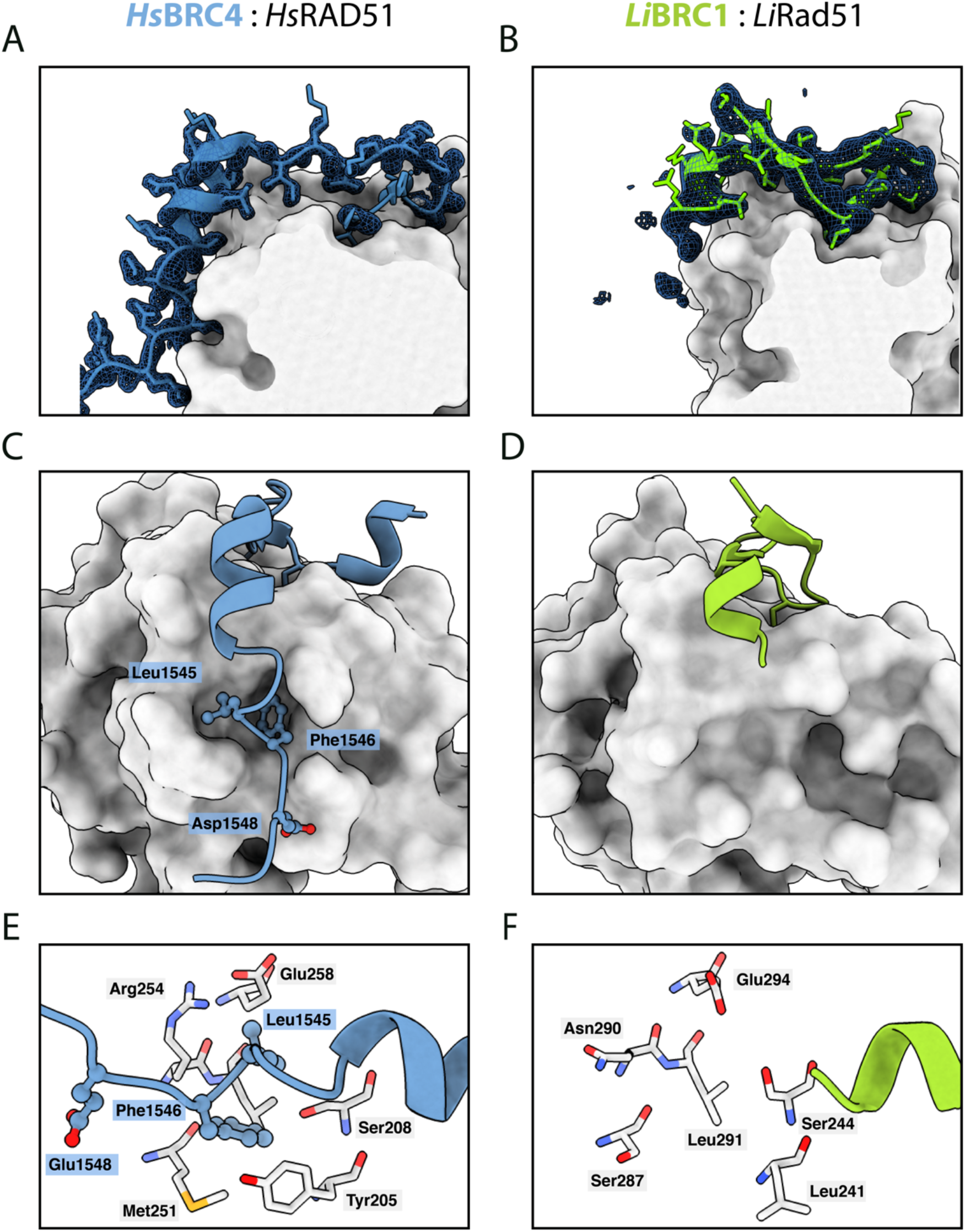
LFDE motif binding is not observed in the *Li*BRC1:*Li*RAD51 crystal structure. (**A**,**B**) 2Fo-Fc electron density maps of (**A**) human BRC4 and (B) *Li*BRC1 at σ = 1. (**C**,**D**) Comparison of the binding modes of C-terminal LFDE motif residues in (**C**) HsBRC4 and (**D**) *Li*BRC1. The difference in hydrophobic pocket depth is clearly apparent. (**E**,**F**) Comparison of the residues involved in the formation of the LFDE-binding cognate hydrophobic pockets in (**E**) human RAD51 and (**F**) *Li*RAD51.

In sum, the crystal structure of the *Li*BRC1:*Li*RAD51 complex reveals a BRC repeat binding-mode defined by an extended β-hairpin at the N-terminus forming a small hydrophobic core, an Arg124-mediated salt-bridge, and a lack of any interaction with *Li*RAD51at the C-terminal LFDE motif.

### *Li*BRC1 C-terminal residues are not involved in binding *Li*RAD51

The X-ray structure prompted us to investigate the contributions to binding of the *Li*BRC1 C-terminus in more detail. A set of *Li*BRC1 mutants were purified and their binding evaluated using the FP competition assay **(Figure 4A)**. Two mutants with extended termini containing additional residues from the full-length protein were evaluated to delineate cut-offs for a binding region, with little change in affinity for the longer repeats observed (*Li*BRC1.2 and *Li*BRC1.3, K_D_ = 0.19 and 0.31 μM, respectively). Step-wise deletions of the C-terminus were then evaluated. Removal of residues up to Arg134 was tolerated without significant loss of affinity (*Li*BRC1.7, K_D_ = 0.39 μM), implying that the 134-RLGD-137 tetrad, whose residue positions correspond to the canonical LFDE motif in humans, is not critical for binding, consistent with our observations from the crystal structure.

**Figure 4.**
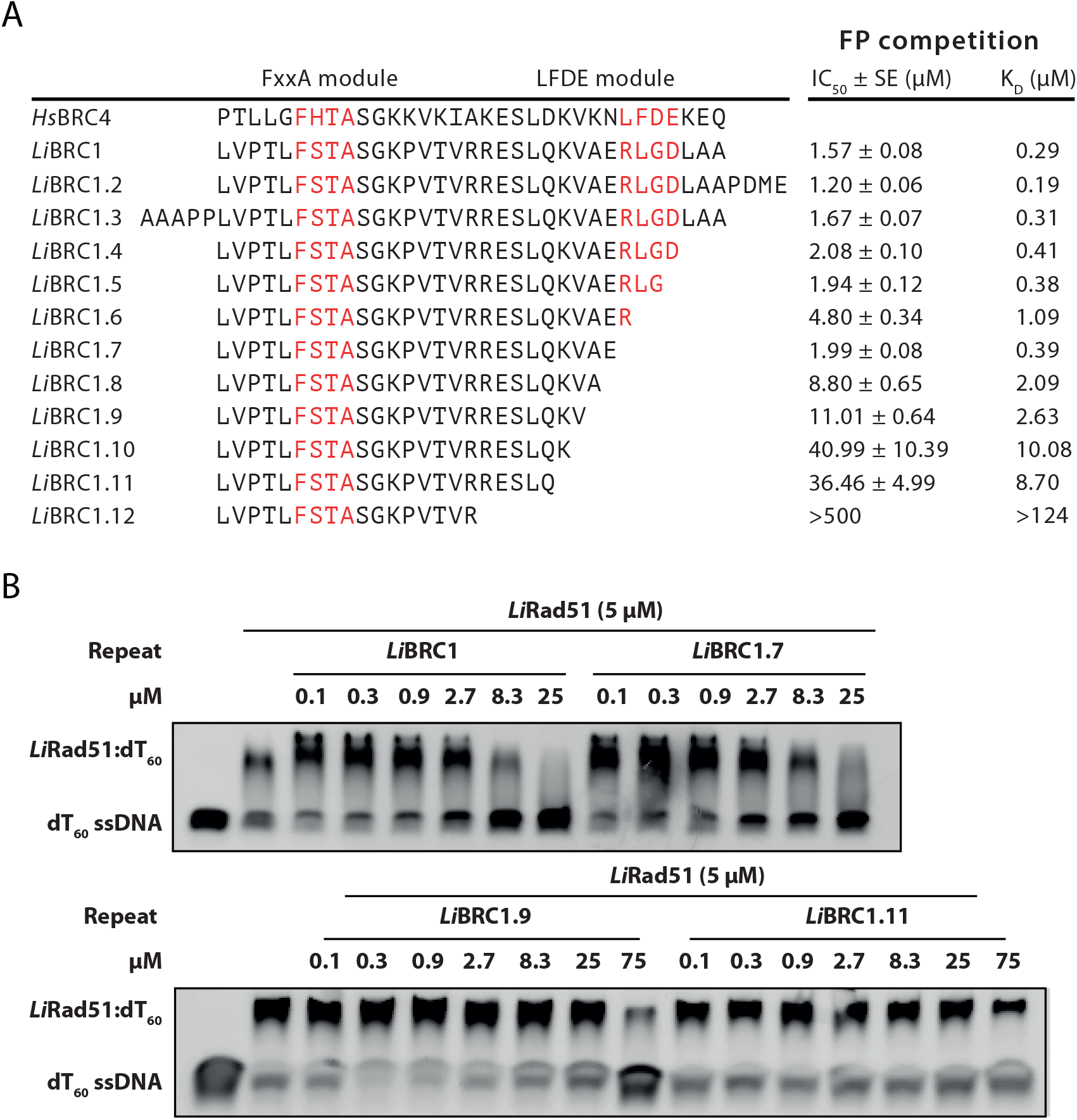
C-terminal LFDE motif residues are not critical for binding *Li*RAD51. (**A**) *Li*BRC1 mutants and their competition FP assay IC_50_ and calculated K_D_ values. SE – standard error of fit. (**B**) Electrophoretic mobility shift assay (EMSA) competition experiments evaluating the ability of *Li*BRC1 truncation mutants to inhibit *Li*RAD51 nucleoprotein filament formation. 5 μM *Li*RAD51 was pre-incubated with GB1-fused *Li*BRC1 or its truncation mutants, after which FAM-labelled ssDNA (dT60, 100 nM) was added to the reaction. Products were resolved on a 1xTBE 2% agarose gel.

The most C-terminal *Li*BRC1 residue observed in the crystal structure is Gln129. Further truncations up until this residue result in a gradual decrease in affinity, reaching a K_D_ of 8.70 μM for *Li*BRC1.11, signifying an important contribution to binding by the 130-KVAE-133 tetrad. Removal of *Li*BRC1 Glu133 causes a more than four-fold drop in K_D_, implying that this residue makes a significant contribution to binding (*Li*BRC1.8, K_D_ = 2.09 μM). It is not immediately apparent from the complex structure how this residue may increase affinity, as there are no nearby *Li*RAD51 side-chains bearing a positive charge to form salt bridges with. It is possible that, rather than interacting with *Li*RAD51, it stabilises the repeat conformation, for example, by interacting with the cationic Lys130 on the same helical face or by affecting the overall charge of the peptide.

To confirm these sequence-activity relationships in the context of the full-length *Li*RAD51 protein, electrophoretic mobility shift assays (EMSA) were performed, in which *Li*BRC1 and its C-terminal truncation constructs *Li*BRC1.7, 1.9 and 1.11 were tested for their ability to inhibit the formation of *Li*RAD51:ssDNA nucleoprotein filament (NF) by competing with *Li*RAD51 self-association (**Figure 4B)**. Both *Li*BRC1 and *Li*BRC1.7 inhibited NF formation in a dose-dependent manner to comparable levels (**Figure 4B**, top). In line with the FP measurements, *Li*BRC1.9 and 1.11 were much less potent inhibitors of NF formation (**Figure 4B**, bottom).

## DISCUSSION

We have shown that the two BRC repeats from the *L. infantum* BRCA2 ortholog bind *Li*RAD51, with *Li*BRC1 manifesting an almost 50 times higher affinity than *Li*BRC2, as measured by an FP competition assay. To our knowledge, this represents the first instance where a direct BRC repeat:RAD51 interaction is confirmed for a non-mammalian BRCA2 ortholog. A number of factors may contribute to the lowered affinity of *Li*BRC2. For example, *Li*BRC2 contains an arginine at position +1 to the FxxA motif, which is occupied by a serine in *Li*BRC1 and most human repeats, forming a hairpin-stabilising hydrogen bond.^19^ Interestingly, human BRC2 also lacks a serine at the equivalent position, and its FxxA module has been shown by repeat shuffling experiments to contribute weakly to RAD51 binding.^27^

A crystal structure of the complex of *Li*BRC1 with the *Li*RAD51 ATPase domain was determined and revealed that *Li*BRC1 residues 134-RLGD-137, corresponding to the conserved LFDE motif in humans, do not form ordered contacts with *Li*RAD51. Subsequent truncation mutagenesis experiments confirmed that, indeed, the LFDE-equivalent part of *Li*BRC1 is not critical for *Li*RAD51 binding. This lack of engagement of LFDE-like motif with RAD51 could explain the lowered affinity of *Li*BRC1 for RAD51 compared to highest affinity human repeats.

Comparative analysis with the human proteins can help rationalise the lack of interaction observed for the 134-RLGD-137 tetrad from the point of view of the repeat sequence, or, alternatively, by looking at the complementary surfaces formed by the ATPase domains. The LFDE motifs of the human BRC repeats are defined by two strongly conserved features. First, two bulky hydrophobic residues, such as Trp, Leu, Phe and Val, are conserved at the first two positions of the motif in all the human repeats. Secondly, an acidic residue at the last position forms a salt-bridge with nearby arginines on human RAD51. The shape of the first two side-chains appears to be less critical than their hydrophobic nature, as evidenced by the different combinations observed in the eight human repeats. Moreover, Rajendra and Venkitaraman showed that exchange of one hydrophobic residue for another is not disruptive for binding, and can in fact bring about improved affinity, such as when Leu1545 is replaced by a tryptophan in human BRC4.^20^ In *Li*BRC1, on the other hand, there is just a single hydrophobic residue present in the amino acid tetrad that corresponds to the LFDE motif, which drastically reduces the buried hydrophobic surface area attainable upon binding. Buried hydrophobic contacts tend to contribute significantly to free energy of binding in protein-protein interactions, therefore it is reasonable to assume that the RLGD tetrad would result in a weaker energetic contribution, even if compensatory contacts, for example, a salt bridge involving Arg134, were present. The cognate surface of the *Li*RAD51 ATPase domain also appears less conducive to the binding of an LFDE-like moiety, as the hydrophobic pockets are less pronounced. In human RAD51, Tyr205 and Met251 form a lining for a deeper LFDE binding site compared to *Li*RAD51.

Remarkably, the *Li*BRC1 repeat manifests sub-micromolar binding in the absence of an LFDE-like interaction. In the crystal structure, the N-terminus of *Li*BRC1 peptide forms an extended β-hairpin, which results in a hydrophobic core folding on the ATPase domain surface, as well as hydrophobic contacts formed by a nearby Leu112. We propose that these additional hydrophobic interactions may partially compensate for the lack of a functional LFDE motif and thus ensure a high affinity interaction for the *L. infantum* BRCA2 ortholog to localise Rad51 to the sites of DNA damage and stimulate nucleofilament formation on resected ssDNA. Further mutagenesis studies will be necessary to deconvolute the contributions of these additional interactions. *Li*BRC1 thus presents a distinct mode of BRC repeat binding for the evolutionary distant *L. infantum*, suggesting that the LFDE motif is not a universal pre-requisite for high-affinity binding, despite previous reports demonstrating that it is indispensable for a functional RAD51:BRCA2 interaction in human cells.^20^

Nucleation of 2-3 RAD51 monomers on ssDNA is the rate-limiting step of RAD51:ssDNA nucleofilament formation and BRCA2 has been proposed to seed RAD51 nuclei on ssDNA.^30,31^ Human BRCA2 can bind up to six RAD51 monomers simultaneously.^15^ While the exact molecular detail of BRCA2-mediated nucleation is not clear, it is likely that high avidity resulting from having more than one BRC repeat may increase nucleation rates. The sequence distance between the two BRC repeats in the *L. infantum* BRCA2 ortholog is much smaller than in human BRCA2, that is, around 6 residues, depending on where repeat boundaries are defined. This means that, in order for the protein to engage more than one RAD51 molecule simultaneously, as has been previously shown for BRCA2, the C-terminus of the *Li*BRC1 repeat may need to be vacant, potentially explaining the lack of interaction for the 134-RLGD-137 tetrad.

Alignment of BRC repeats from a set of representative eukaryotes indicates that other protozoans may have similarly divergent binding modes (**Figure S3**). For example, BRC repeats from both *Trypanosoma* and *P. falciparum* contain a threonine at -2 and a valine/leucine at -4 to the FxxA motif, which suggests formation of a similar extended β-hairpin and a hydrophobic core by the repeat. Moreover, the repeats from these same organisms lack a human-like LFDE motif. It is thus possible that the evolution of BRC repeats was first defined by the formation of a universally conserved FxxA motif in a common ancestor, which closely mimics the RAD51 self-oligomerisation interface, and was then followed by subsequent steps of affinity fine-tuning in which different additional features evolved for distinct eukaryote clades.

## AVAILABILITY

All data are available from the authors upon request.

## ACCESSION NUMBERS

Atomic coordinates and structure factors for the reported crystal structures have been deposited with the Protein Data bank under accession number 7QV8.

## ACKNOWLEDGEMENT

We thank Diamond Light Source (Harwell, UK) for the access to the i04-1 beamline. We are grateful for support and access to instrumentation at the X-ray crystallographic facility and the biophysics facility at the Department of Biochemistry, University of Cambridge. We thank Dr Joseph Maman for his advice on setting up the fluorescence polarisation assay.

## SUPPLEMENTARY DATA

**Table S1.** Crystallisation conditions, data collection and refinement statistics. Values in parentheses are for the high-resolution bin.

|  |  |
| --- | --- |
| **Crystallisation** |  |
| Protein | 0.5 mM *Li*BRC1 :*Li*RAD51<sup>ATPase,ΔL2</sup> in 20 mM Tris pH 8.0, 100 mM NaCl, 100 mM Li<sub>2</sub>SO<sub>4</sub>, 20 mM ADP/MgCl<sub>2</sub> |
| Crystallisation solution | 32% low MW PEG smear (precipitant, Molecular Dimensions), 0.1M Tris pH 8.5 |
| Protein:solution volume (nl:nl) | 200:200 |
| PDB | 7QV8 |
| **Data collection processing** |  |
| Beamline | DLS i04-1 |
| Wavelength (Å) | 0.9159 |
| Space group | P 4 2 2 |
| a, b, c (Å) | 61.00 61.00 119.22 |
| α, β, γ (°) | 90.00 90.00 90.00 |
| Resolution range (Å) | 2.15 - 59.61 (2.15 - 2.18) |
| R<sub>meas</sub> | 0.119 (8.167) |
| Completeness (%) | 98.9 (98.9) |
| Number of total / unique reflections | 296871 / 12826 |
| Redundancy | 23.1 (18.1) |
| <I/σ(I)> | 15.6 (0.4) |
| CC<sub>1/2</sub> | 1.0 (0.5) |
| **Refinement** |  |
| R<sub>cryst</sub> / R<sub>free</sub> | 0.211 / 0.267 |
| Resolution range (Å) | 54.31 - 2.15 |
| Number of reflections: work/test set | 10597 / 517 |
| Number of atoms | 1806 |
| Mean/Wilson B-factor | 67.897 / 50.7 |
| Ramachandran<br>favoured/allowed/outliers (%) | 98.65 / 1.35 / 0.00 |
| RMSD bonds (Å) | 0.012 |
| RMSD angles (°) | 1.62 |

**Table S2.**
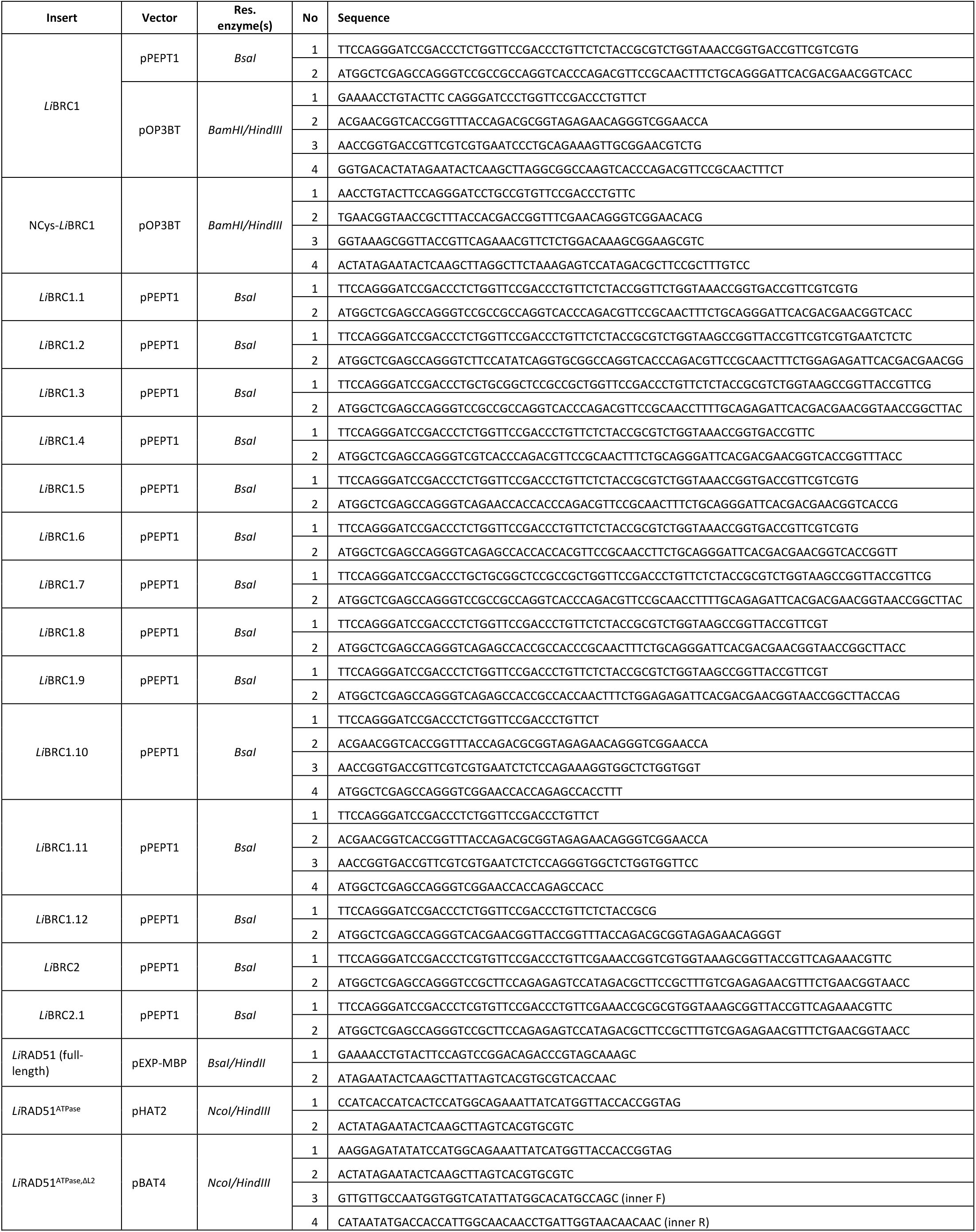
Oligonucleotides used for assembly PCR and cloning.

**Figure S1.**
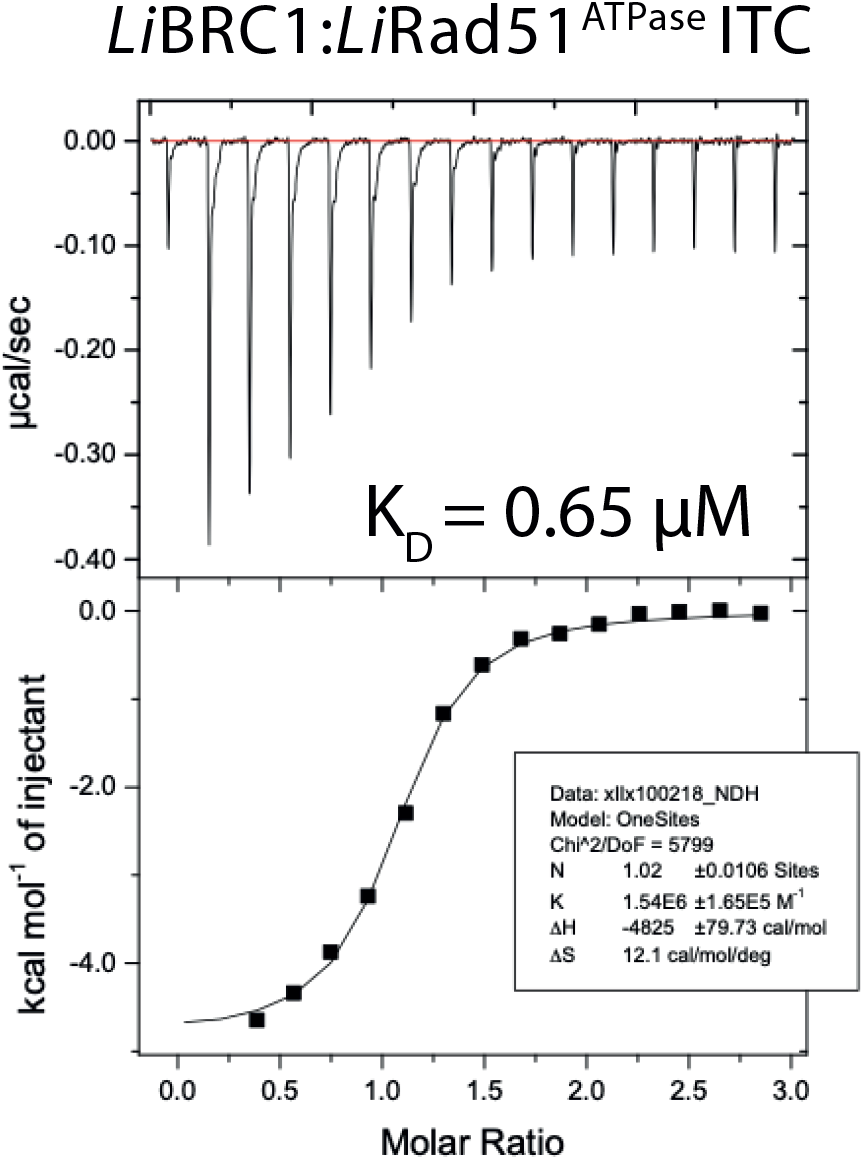
ITC titration of *Li*BRC1 peptide into *Li*RAD51^ATPase^.

**Figure S2.**
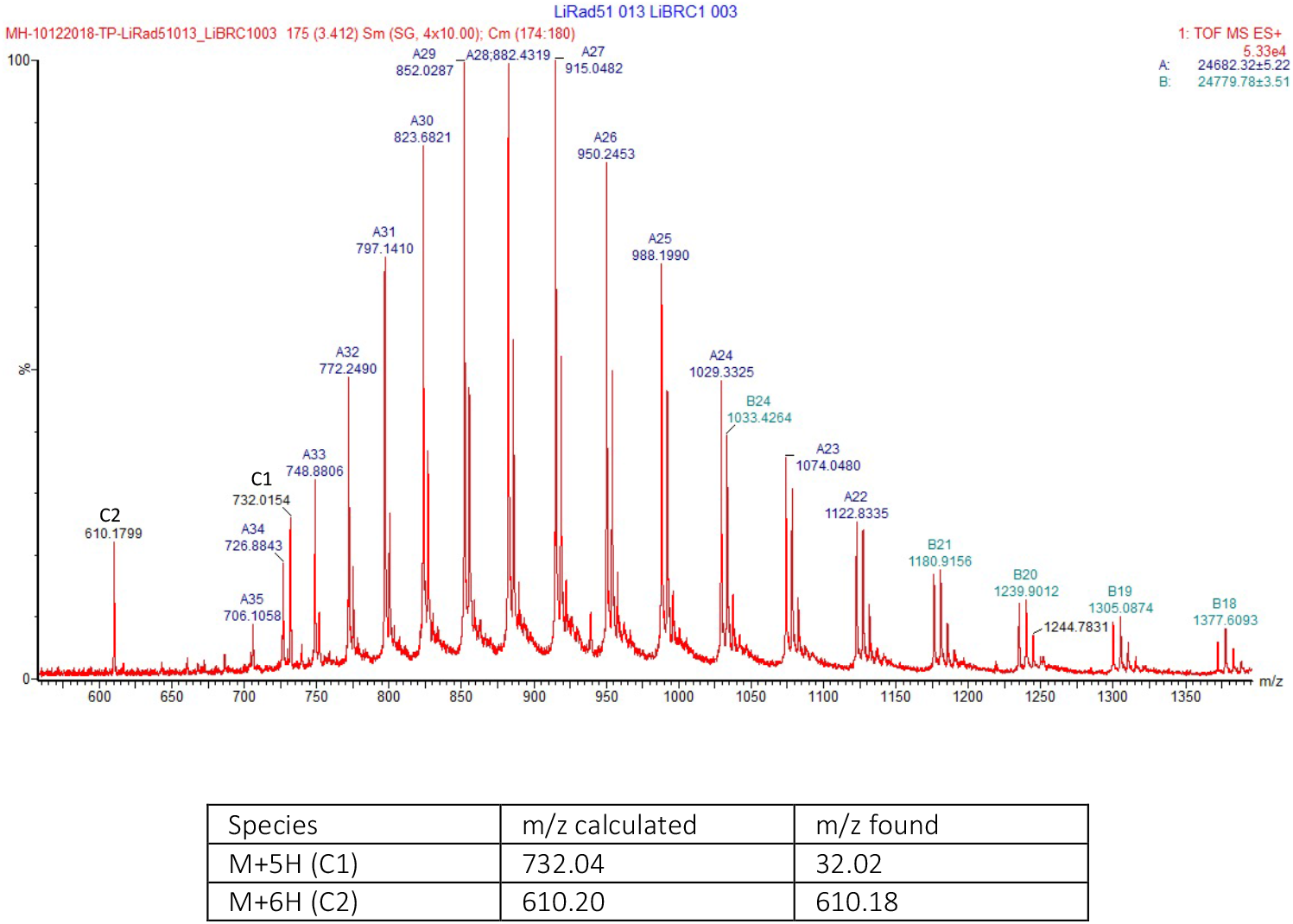
Protein mass spectrum of the *Li*BRC1:*Li*RAD51^ATPase,ΔL2^ complex. Peaks C1 and C2 correspond to the full-length *Li*BRC1 peptide.

**Figure S3.**
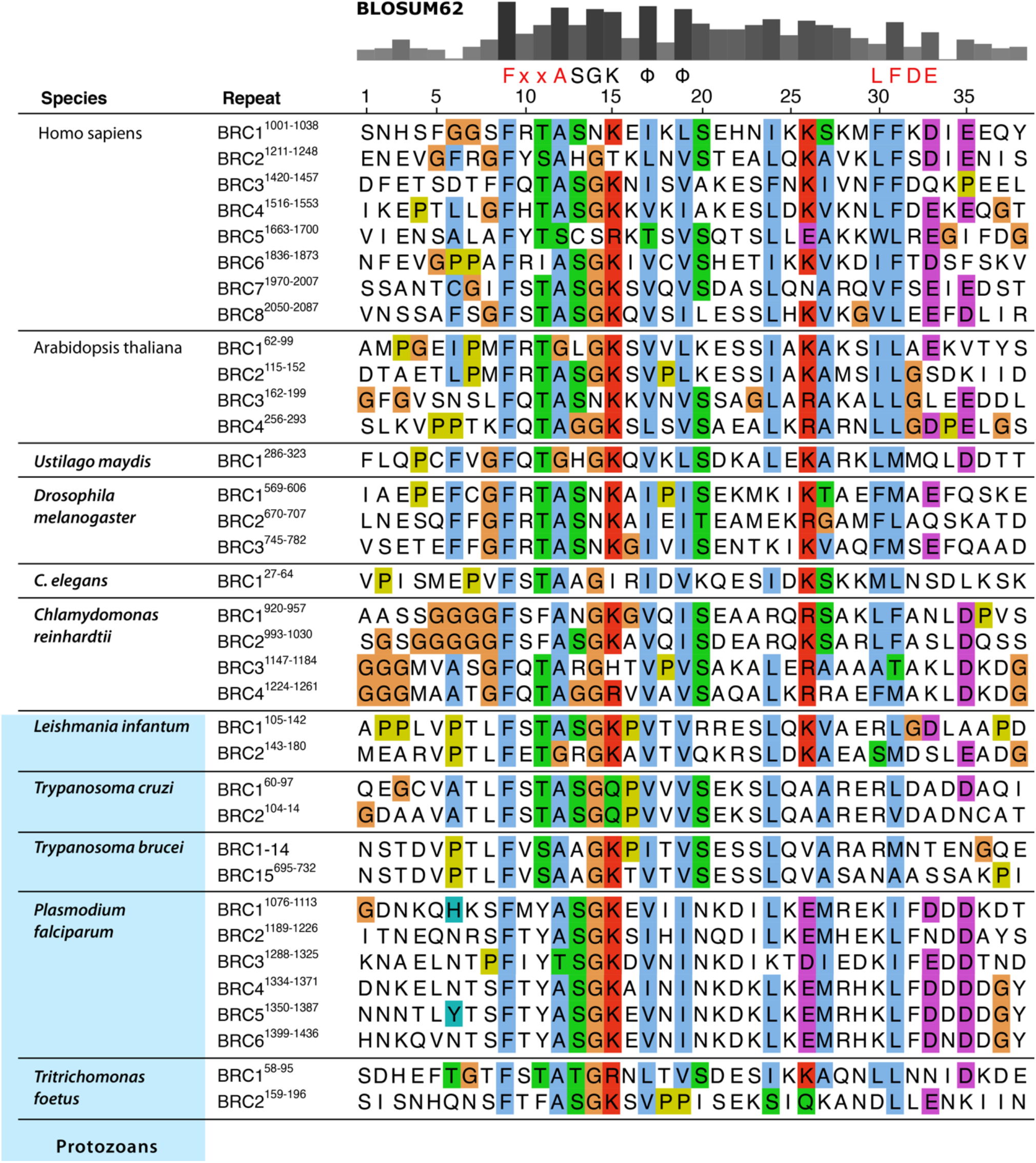
Sequence alignment of BRC repeats from a set of representative organisms and protozoan parasites. The grey bars on top represent BLOSUM62 alignment scores. Conserved residues are coloured using the default ClustalX colour scheme.

**Figure S4.**
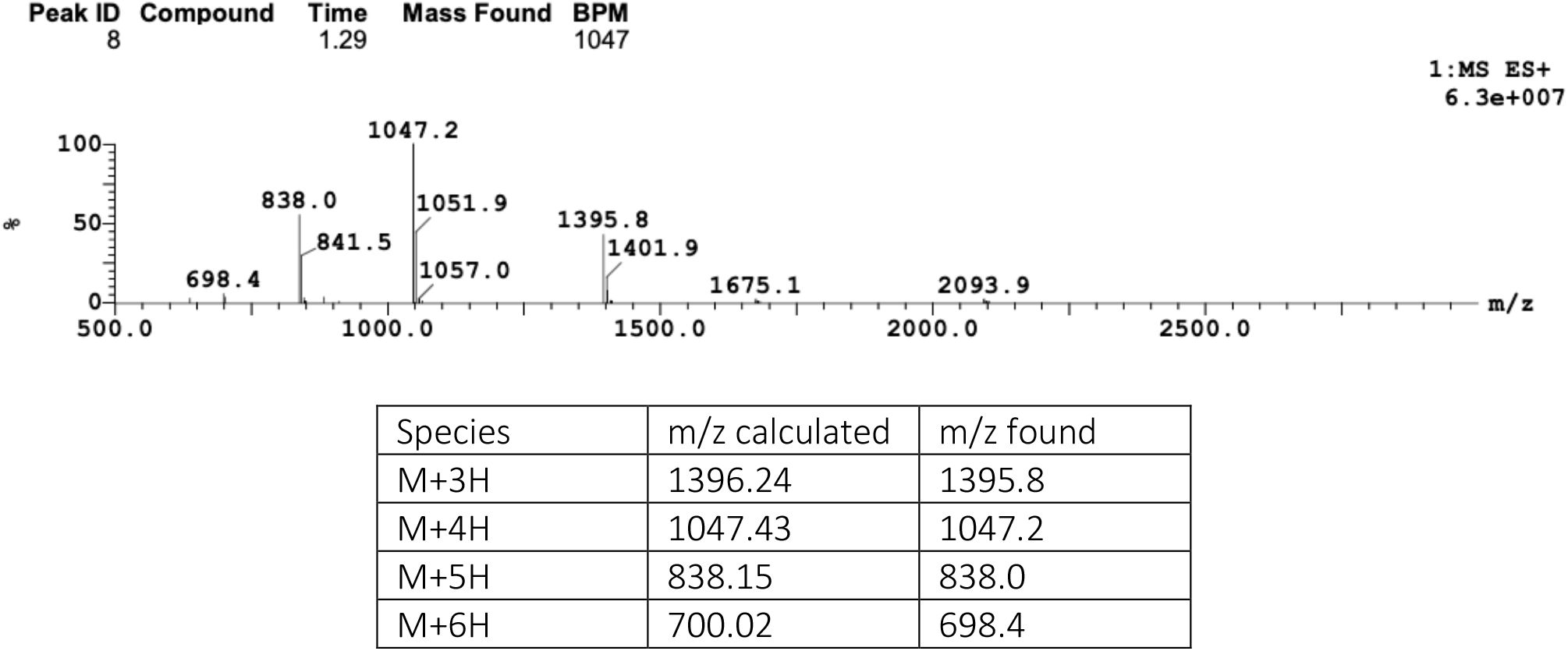
Mass spectrum of the *Li*BRC1-fluor peptide.

